# Using fecal immunochemical tubes for the analysis of the gut microbiome has the potential to improve colorectal cancer screening

**DOI:** 10.1101/2021.03.15.435399

**Authors:** Kertu Liis Krigul, Oliver Aasmets, Kreete Lüll, Tõnis Org, Elin Org

**Affiliations:** Institute of Genomics, Estonian Genome Centre, University of Tartu, Tartu, Estonia; Department of Biotechnology, Institute of Molecular and Cell Biology, University of Tartu, Tartu, Estonia

**Keywords:** Colorectal cancer, Gut microbiome, Sample preservation, Fecal immunological test

## Abstract

**Background:** Colorectal cancer (CRC) is an important and challenging public health problem which successful treatment depends on the early detection of the disease. Recently, colorectal cancer specific microbiome signatures have been proposed as an additional marker for CRC detection. A desirable aim would be the possibility to analyze microbiome from the fecal samples collected during CRC screening programs into FIT tubes for fecal occult blood testing.

**Methods:** We investigated the impact of the Fecal Immunohistochemical Test (FIT) and stabilization buffer on the microbial community structure in stool samples from 30 volunteers and compared their communities to fresh-frozen samples highlighting also the previously published cancer-specific communities. Altogether 214 samples were analyzed including positive and negative controls using 16S rRNA gene sequencing.

**Results:** The variation between individuals is greater than the differences introduced by collection strategy. The vast majority of the genera are stable for up to 7 days. None of the changes observed between fresh frozen samples and FIT tubes are related to previously shown colorectal-cancer specific bacteria.

**Conclusions:** Overall, our results show that FIT tubes can be used for profiling the gut microbiota in colorectal cancer screening programs as the community is similar to fresh frozen samples and stable at least for 7 days.

**Impact:** Sample material from FIT tubes could be used in addition to fecal immunochemical tests for future investigations into the role of gut microbiota in colorectal cancer screening programs circumventing the need to collect additional samples and possibly improving the sensitivity of FIT.

## INTRODUCTION

Colorectal cancer (CRC) affects millions of people worldwide each year and is one of the leading causes of death among cancers. Due to its high frequency, CRC has become an important and challenging public health problem where the detection of cancer at its early stages is of high importance. Therefore, many countries all over Europe and the world have started population-based screening programs, which aim to detect CRC by analyzing fecal blood using immunochemical fecal occult blood test (iFOBT/FIT) followed by invasive colonoscopy.

Screening programs face multiple challenges. Depending on the country, the screening programs invite individuals in the age range of 50-74 to participate (1). Recent data shows, however, that the incidence of colorectal cancer is increasing especially among younger adults (2). On average, half of the invited patients do not participate in FIT-based screening programs, even though there is more than a 90% chance of survival when the cancer is detected in early stage (1,3). Furthermore, commonly used FIT tests have shown low sensitivity for colorectal cancer lesions (sensitivity for non-advanced adenomas is 7.6 %) (4). Additionally, FIT tests could have false negative results due to smoking and advanced age, both well-known risk factors for colorectal cancer (5), resulting in additional colorectal cancer cases left unnoticed. Moreover, the data has shown that over 20% of colorectal adenomas can be missed in colonoscopy which is considered the golden standard of colorectal cancer diagnosis (6,7). In addition, around 30% of FIT-positive individuals undergoing colonoscopy might have negative colonoscopy (i.e. normal colon without any pathologies) (8). Due to the aforementioned reasons, highly specific, inexpensive, and sensitive non-invasive screening tests and additional biomarkers to increase sensitivity are urgently needed. Gut microbiome has been proposed as a potential additional biomarker.

Recent studies indicate that the gut microbiome plays an important role in development of the immune system (9), etiology of metabolic (10) as well as neurological diseases (11), and cancer (12–14). The cross-sectional multi-population human studies have shown significant associations between gut microbiota and colorectal cancer where colorectal cancer microbiome (14) as well as cancer-stage specific microbial signatures (15) have been detected from stool samples of CRC patients. It has been suggested that complementing fecal occult blood test with gut microbiota could improve detection of colorectal cancer compared to using the fecal occult blood test alone (3,16). As both tests use fecal samples, it would be favorable to obtain both fecal occult blood test and microbiome composition results from the same sample.

It has been demonstrated recently that fecal immunochemical test (FIT) tubes used for fecal occult blood sample collection have the potential to be also used as a sample collection method for microbiome studies (17,18). An extensive amount of FIT tubes is available on the market that have different composition and have been shown to perform differently detecting fecal occult blood from colorectal cancer patients (19), indicating the possibility that they might also vary in performance to detect the microbiome. The Colorectal Cancer Screening in multiple European countries (e.g. Sweden, Finland, Estonia, Czech Republic, and Slovakia) use QuikRead^®^ iFOB Sampling Set (Aidian, Espoo, Finland), however, these particular FIT tubes have not yet been tested as a method for microbiome sample collection and analysis. Using fecal samples not originally intended for microbial profiling may introduce technical challenges due to incompatible materials and varying sample handling and storage conditions. Therefore, the aspects of how sample processing and storage may potentially influence the microbiota need to be investigated.

In the current study, we aimed to explore the potential to determine the composition of the gut microbiome from the FIT tubes of the QuikRead^®^ iFOB Sampling Set. To test this, we analyzed 30 volunteers’ fecal samples stored in FIT tubes and compared them with two more storage methods - fresh-frozen or storage in stabilizing solution DNA/RNA Shield (Zymo Research, Irvine, California). We investigated how storage in FIT and stabilizing solution DNA/RNA shield affect gut microbiome diversity and composition estimates as well as the community structure stability over time, compared to the gold standard of immediate freezing.

## MATERIALS AND METHODS

### Sample population and collection

30 volunteers were recruited in the study, who contributed their fecal samples for microbiome analyses. 16 (53.3%) of the participants were female and 14 (46.7%) were male. All recruited subjects were Estonians aged between 22 and 68 (39 ± 12.1) with BMI ranging from 18.4 to 41.8 kg/m2 (mean 24 ± 4.7 kg/m^2^). Written consent was obtained from the volunteers and the study followed the sampling protocols which were approved by the Ethics Committee of the University of Tartu.

Seven samples were collected for each volunteer and in total 214 samples (including positive and negative controls) were analyzed in this study. All the samples from the volunteers were collected within the same week (January, 2020). A fresh stool sample was collected immediately after defecation with a sterile Pasteur pipette and placed inside a polypropylene conical 15 ml tube without a preservative and then frozen at −20°C (the “gold standard” of microbiome studies). From the same fecal sample, each individual collected 3 subsamples in a QuikRead go iFOBT fecal immunochemical test tube (Aidian, Espoo, Finland) using a stick attached to the lid according to the instructions provided in the kit. Additionally, 3 aliquots were collected in 1 ml stabilization buffer tubes (DNA/RNA Shield, Zymo Research, Irvine, California) using sterile swabs. In order to assess how storage time affects the stability of the community structure, one FIT tube and one stabilization buffer tube were frozen instantly together with the fresh stool sample. These tubes were frozen within 16 minutes (SD+16.9) after the sample was taken. Additional FIT and stabilization buffer tubes were stored at room temperature either 96 h (4 days) or 168h (7 days) and then frozen at −20 °C. The rationale for doing this was that in CRC screening programs, the time after collecting the initial fecal sample until arriving at the study center for occult blood testing can take up to a week. Therefore, we wanted to see if longer shipping times could compromise the stability of the microbial community and affect the results of microbiome analysis. Furthermore, in addition to the ZymoBIOMICS^™^ Microbial Community Standard or MOCK (Zymo Research, Irvine, California) used as a positive control for sequencing, negative controls for each sample type were used for DNA extraction and sequencing steps. Workflow for the sample collection and storage conditions is shown in Figure 1.

**Figure 1.**
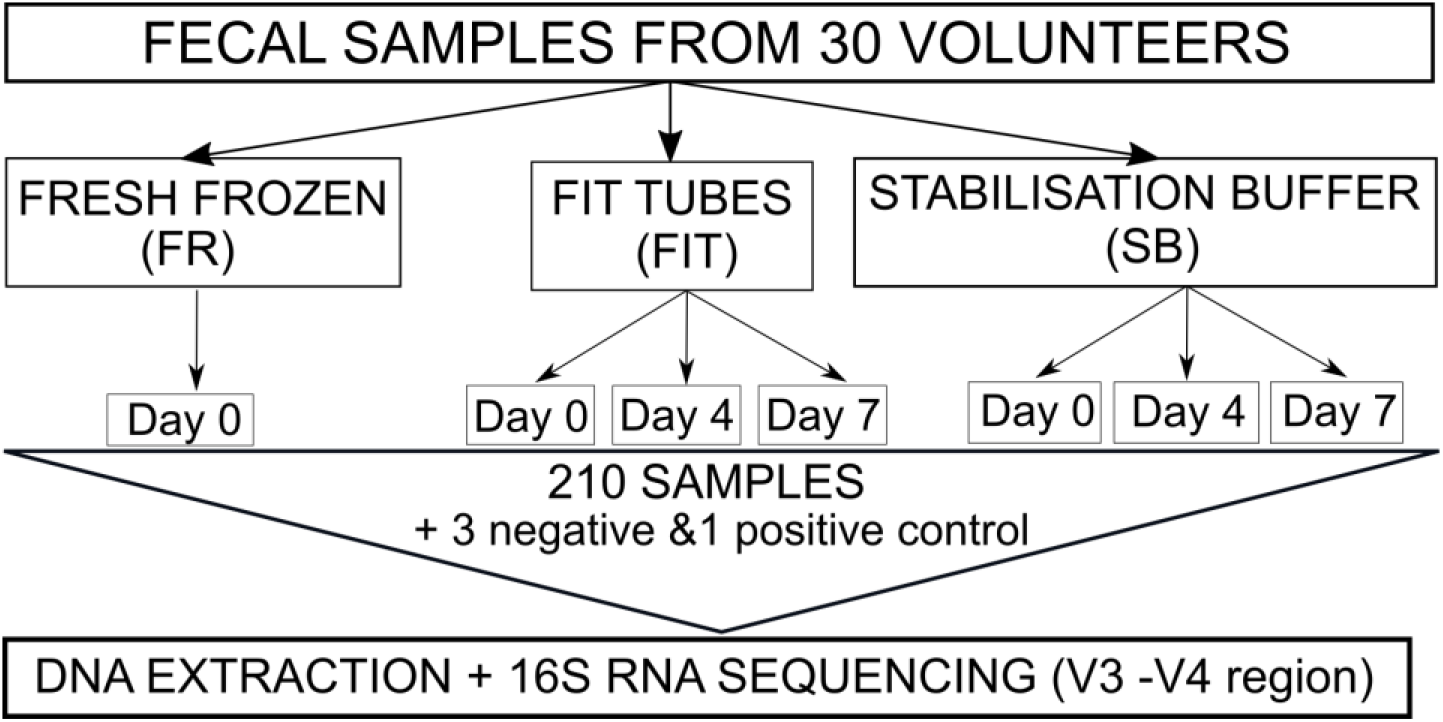
Workflow for sample collection and analysis.

### DNA extraction and sequencing

DNA extraction for all samples was done using Qiagen DNeasy PowerSoil Pro DNA extraction kit (Qiagen, Venlo, The Netherlands). For fresh-frozen samples, around 200 mg of stool was used as a starting material following the DNA extraction kit manufacturer’s instructions with the exception that the samples were incubated additional 10 minutes at 65 °C after adding solution CD1 to ensure proper lysis of difficult to lyse bacterial cells. The cell disruption step was done using Precellys 24 tissue homogenizer (parameters: 2×30 sec, 2500 rpm, 30 second break) (Bertin Instruments, Montigny-le-Bretonneux, France). For the samples stored in the stabilization buffer, 250 μL of liquid was used as a starting material. The rest of the protocol was the same as with fresh-frozen samples. For the samples in the FIT tubes, up to 2 ml of the FIT solution was transferred into a new tube and 200 μL of 1M Tris-HCl, pH 7.5 was added to quench formaldehyde present in the FIT solution. After centrifugation, the supernatant was discarded, the pellet was taken up in CD1 solution and added to the PowerSoil Pro tubes. To increase DNA yield, decrosslinking was performed by 4-hour incubation at 65°C with Proteinase K before the cell disruption step. The rest of the protocol was done following the manufacturer’s instructions. DNA was quantified from all samples using Qubit Fluorometer using dsDNA HS Assay Kit (Thermo Fisher Scientific) and diluted to 5 ng/μL for sequencing. DNA extraction protocol was also followed through using negative controls (no solution, as well as FIT and stabilization buffer tubes with their original solution).

### Microbial community analysis

The amplicon sequencing was conducted as follows in the Institute of Genomics Core Facility, University of Tartu. Extracted DNA samples were quantified with Qubit^®^ 2.0 Fluorometer (Invitrogen, Grand Island, USA). The genomic DNA was amplified using primers 16S_F (5′- TCGTCGGCAGCGTCAGATGTGTATAAGAGACAGCCTACGGGNGGCWGCAG −3′) and 16S_R (5′-GTCTCGTGGGCTCGGAGATG TGTATAAGAGACAGGACTACHVGGGTATCTAATCC −3′) for PCR amplification of an approximately 460 bp region within the hypervariable (V3-V4) region of prokaryotic 16S ribosomal RNA gene (20). Amplicon libraries for Illumina (Illumina, San Diego, USA) nextgeneration sequencing were generated by two-step PCR. First, region specific for 16S rRNA was amplified with 24 cycles and then Illumina adapter and index sequences were added by 7 cycles of PCR. The quality control of amplicon libraries was performed by Agilent 2200 TapeStation analysis (Agilent Technologies, Santa Clara, USA) and with Kapa Library Quantification Kit (Kapa Biosystems, Woburn, USA). Amplicon libraries were pooled in equimolar concentrations. Sequencing was carried out on an Illumina MiSeq System using MiSeq Reagent Kit v3 in paired end 2 × 300 bp mode.

Raw sequences were demultiplexed with Illumina bcl2fastq2 Conversion Software v2.20. Bioinformatics analyses were performed using open-source software QIIME2 (version 2019.7) (21). Raw data was imported using the q2-tools import script with PairedEndFastqManifestPhred33 input format. In total, 7,468,645 reads were generated (on average 34 738 reads per sample). The total number of reads for fresh samples (FR) was 1,116,409 (on average of 37 214 reads), for FIT Day 0 (FIT0) samples 1,048,723 reads (on average of 34 957 reads), for FIT Day 4 (FIT4) 1,066,370 reads (on average of 35 546 reads), and for FIT Day 7 (FIT7) 1,066,370 reads (on average of 34 222 reads). For the stabilization buffer DNA/RNA Shield the number of reads for Day 0 (SB0) was 1,028,579 reads (on average of 34 286 reads), for stabilization buffer Day 4 (SB4) 1,127,694 reads (on average 37 590 reads), and for stabilization buffer Day 7 (SB7) 1,026,705 reads (on average of 34 224 reads).

Denoising was done using DADA2 software, which uses a quality-aware model of Illumina amplicon errors to attain an abundance distribution of sequence variance, which has a difference of a single nucleotide (22). Based on the quality scores, q2-dada2-denoise script was used to truncate the forward reads at position 250 and trimmed at position 15, and reverse reads were truncated at position 247 and trimmed at position 12. The chimeras were removed using the “consensus” filter which detects the chimeras in each sample individually. With this method, the sequences which are established as chimeric in a fraction of samples are removed. During the denoising steps, forward and reverse reads are also merged. Subsequently, aligning of Amplicon Sequence Variants (ASVs) was done using MAFFT (23). Thereafter, FASTTREE was used to construct phylogeny (24). The taxonomy was assigned using the q2-feature-classifier with the pre-trained naïve Bayes classifier, which was based on the reference reads from SILVA 16S V3-V4 v132_99 databases with similarity threshold of 99% (25,26). All samples passed the quality control (QC) and negative controls, as expected, resulted in 0 ASVs after quality control steps.

### Statistical analysis

Statistical analyses were carried out in RStudio (version 1.2.1335, R version 3.6.1) using packages *phyloseq* (v.1.30.0) (27), *microbiome* (v.1.8.0) (28), *vegan* (v.2.5-6) (29), stats (v.3.2.1) and *ALDEx2* (v.1.18.0) (30). All the visualizations were made using the *ggplot2* (v.3.2.1) (31). Alpha diversity metrics such as the number of genera (estimates the richness of the sample) and Shannon diversity Index (takes into account both samples’ richness and evenness) were calculated on the genus-level microbiome profile using the *phyloseq* package. Between-sample distances were calculated using Euclidean distance metric on centered log ratio (CLR) transformed genus-level microbiome profile (32). Permutational Analysis of Variance (PERMANOVA) on between-sample distances was carried out to test whether differences in microbial composition (beta-diversity) are associated with sample type and time sample spent on room temperature. PERMANOVA was done using *adonis* function from *vegan* package (v.2.5-6.). *Microbiome* package (v.1.6.0) was used to determine the core genera of the microbiome with a detection threshold of 0 and prevalence threshold of 95%. Welch’s paired t test integrated in the ANOVA-Like Differential Expression tool (ALDEx2, v.1.18.0) was used for differential abundance analysis of genera to assess whether FIT or stabilization buffer samples differ from fresh-frozen samples (accuracy of different collection strategies) as well as if there are differences between samples frozen immediately compared to samples frozen on day 4 or day 7 (stability over time). In order to limit the number of tests, the genera whose prevalence was less than 10% were filtered out, leaving 171 out of 360 genera for the analysis. Multiple testing was taken into account using the Benjamini-Hochberg False Discovery Rate (FDR) method and p-values < 0.05 were considered to be statistically significant (33).

## RESULTS

### Study design

For each individual in the study, seven stool samples were collected and stored using 3 different methods prior DNA extraction: 1) fresh, immediately frozen stool samples (FR samples); 2) stool stored in FIT tubes: immediately frozen (FIT0), stored 4 days (FIT4) or 7 days (FIT7) at room temperature; 3) stool stored in DNA/RNA shield stabilization buffer: immediately frozen (SB0), stored 4 days (SB4) or 7 days (SB7) at room temperature) (Figure 1). As expected, the highest DNA concentration was obtained with the fresh-frozen samples (mean 374,23 ng/ul), followed by stabilization buffer samples (mean 42,64 ng/ul) and FIT samples (mean 11,68 ng/ul) (Supplementary Table 1). For each sample, we generated amplicon libraries targeting the bacterial 16S rRNA V3-V4 region, sequenced, performed quality filtering and ASV estimation in QIIME (see Methods). Following the QC step, the average number of reads per sample remained relatively stable across all the sample types (Supplementary Table 1). Negative controls had no read counts after QC step. MOCK community, which was used as a positive control for sequencing, had 18 890 reads after QC step. All genera expected to be present in positive controls (*Staphylococcus, Pseudomonas, Enterococcus, Escherichia, Salmonella, Lactobacillus, Listeria*, and *Bacillus*), were detected in the analysis (Supplementary Table 2). In total, we detected 11 948 ASVs, 360 genera, 131 families, 64 orders, 32 classes, and 18 phyla. At day 0, all the sample types captured similar taxonomic profiles in both phylum and genus taxonomic level (Figure 2, Supplementary Table 3). As expected, a Western microbial community structure was observed in all of the sample types with 90% of bacteria belonging to the phyla *Firmicutes* and *Bacteroides*, which are followed by phyla *Proteobacteria, Actinobacteria*, and *Verrucomicrobia* (Figure 2A).

**Figure 2.**
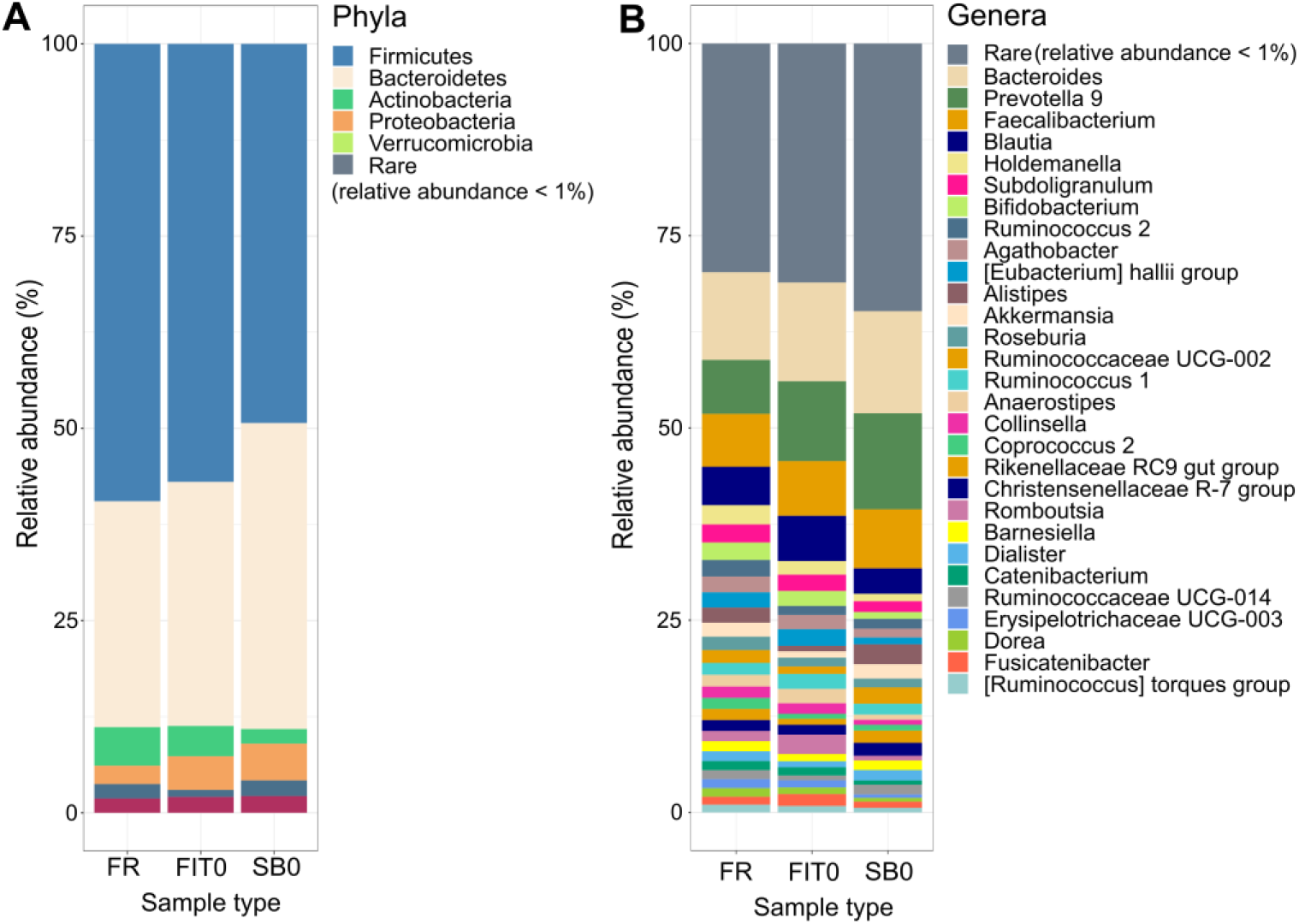
Relative abundance of phyla (A) and genera (B) in different collection methods on day 0. Taxa with mean relative abundance less than 1% are grouped into rare category. Abbreviations: FR – fresh-frozen; FIT0 – immediately frozen FIT samples; SB0 – immediately frozen stabilization buffer samples.

### Impact of sample collection strategy on the diversity of gut microbiome

To evaluate if FIT and stabilization buffer samples have similar diversity to fresh-frozen samples, we assessed differences in the gut microbiome alpha and beta diversity between the collection methods. Both alpha diversity (richness and Shannon index) and beta diversity metrics were calculated using genus-level transformed data. We detected 98 ± 17,6 genera in FR samples, 96 ± 16,3 genera in FIT0, and 95 ± 17,4 in SB0 samples (Supplementary table 4). When comparing richness among the sample types, we found that the differences between the observed genera were not significant between fresh and the other two sample types (FDR_FR-FIT0_ = 0.12, FDR_FR-SB0_ = 0.086) (Figure 3A, Supplementary table 4). However, the samples stored in FIT and stabilization buffer exhibited lower Shannon index values relative to fresh frozen samples (FR_Shannon_ = 3.5 ± 0.3, FIT0_Shannon_ = 3.4 ± 0.3, SB0_Shannon_ = 3.3 ± 0.3, paired t test FDR < 0.01) (Figure 3B, Supplementary table 4). This trend was not only observable between mean Shannon index values in each storage condition, but was also noticeable when each individual’s samples were visualized. When evaluating beta diversity which represents how much the community changes between sample types, we saw that the samples of the same individual group together regardless of storage conditions, indicating that differences between subjects are greater than differences between storage methods (Figure 3C). Analyzing the significance of variance (PERMANOVA) in samples frozen immediately after collection, we found sample type to be significant, however the effect on variance was low (R^2^ = 0.2226, p < 0.001).

**Figure 3.**
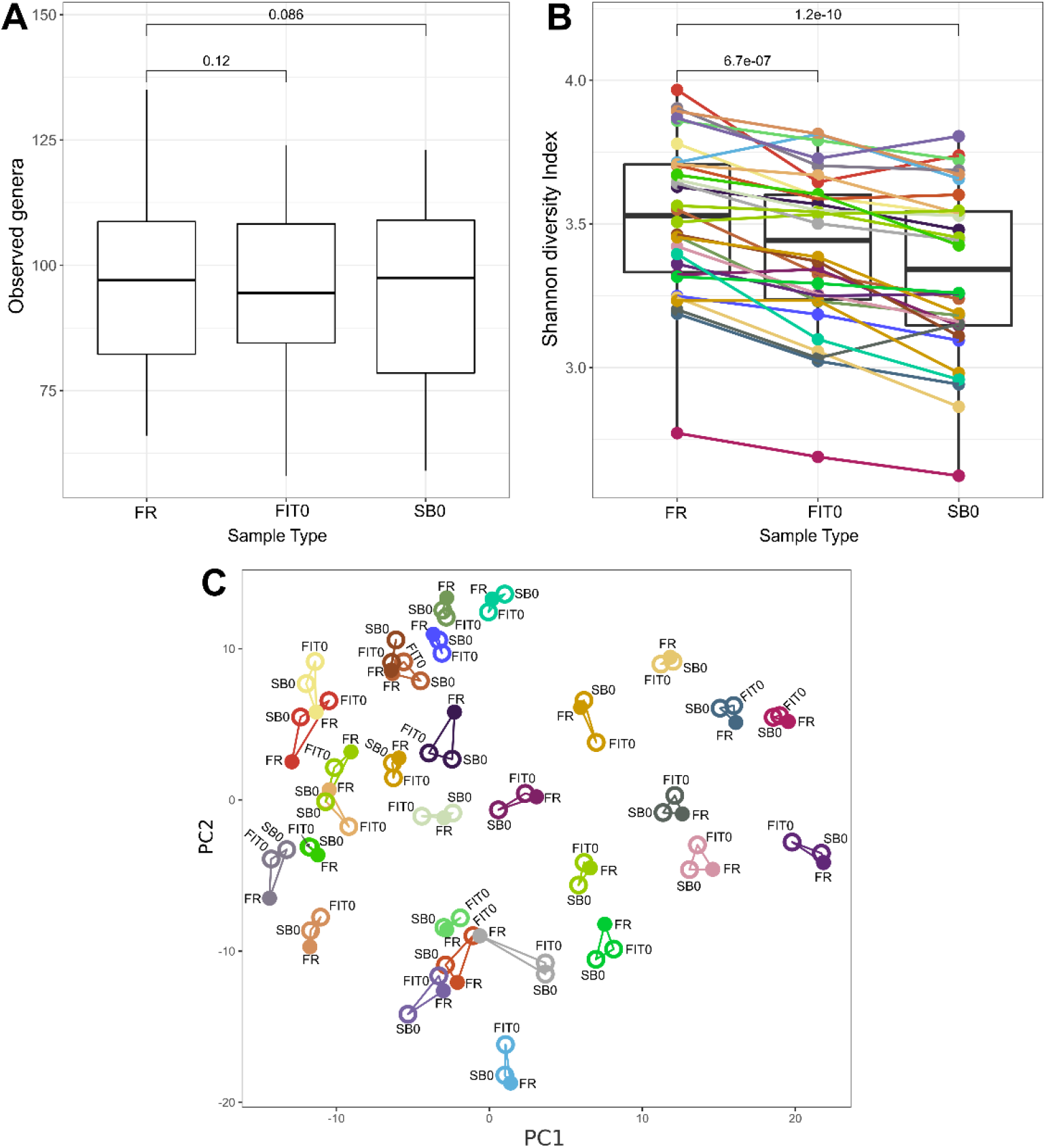
Comparison of microbiome diversity between different sample types. Boxplots represent two different alpha diversity measurements: (A) richness or the number of taxa observed and (B) Shannon diversity index. Median values and interquartile ranges have been indicated in the plots. In richness analysis, paired t tests indicated that the differences are not significant (FDR_FR-FIT0_ = 0.12, FDR_FR-SB0_= 0.086). In Shannon diversity index, samples from the same patient are connected and colored to illustrate the lower trend of alpha diversity for FIT0 and SB0 compared to FR (FDR < 0.01). (C) Principal component analysis (PCA) of beta diversity is shown between storage conditions. Samples are colored and linked based on the individual’s ID. Abbreviations: FR – fresh-frozen; FIT0 – immediately frozen FIT samples; SB0 – immediately frozen stabilization buffer samples.

### Differentially abundant genera between sample collection strategies

In order to test the differences between genera abundance, we used ALDEx2 to identify differentially abundant genera between fresh frozen samples and FIT and stabilization buffer samples. The genera whose prevalence in all of the samples was less than 10% were filtered out, leaving 171 out of 360 genera for the analysis. Out of 171 genera analyzed, we observed 7 genera (4%) with statistically different abundance between FR and FIT0 samples and 16 genera (9,4%) with different abundance between FR and SB0 samples (Figure 4, Supplementary table 5). The rest of the genera abundances were not significant after correction for multiple testing (FDR > 0.05). Six genera with significantly different abundance between FIT0 and FR samples (*Alistipes, Anaerostipes, Eubacterium coprostalinogenes* group, *Romboutsia*, and uncultured *Ruminococcaceae*) had also significantly different abundance in SB0 samples when compared to FR samples, but only in the *Eubacterium coprostanoligenes* group the change was in the same direction for both FIT0 and SB0.

**Figure 4.**
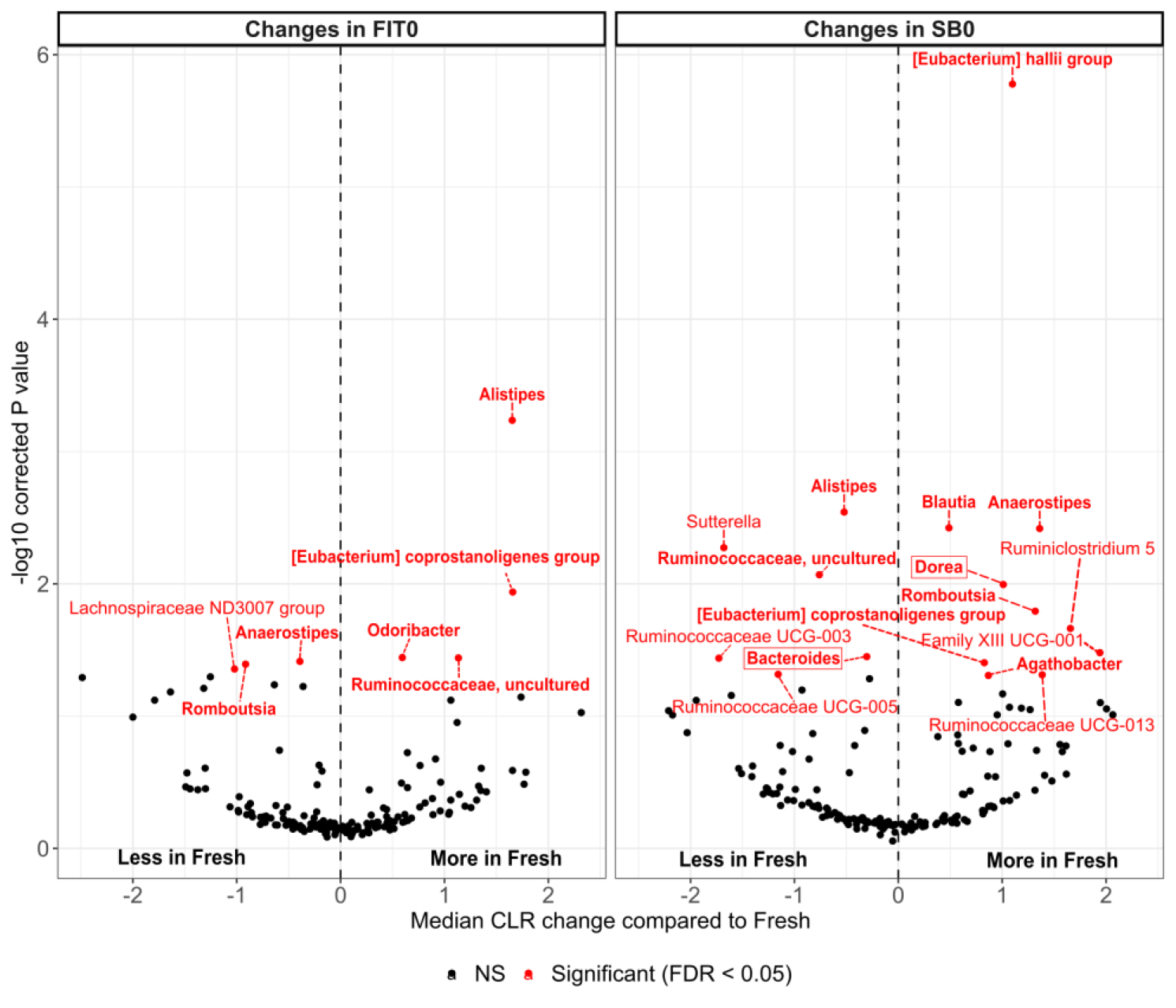
Differentially abundant genera between different sample types. Average CLR changes in FIT0 and SB0 compared to FRESH samples are shown where significantly different taxa (Benjamini-Hochberg correction, FDR < 0.05) are colored in red. The genera belonging to the core 95% are indicated in bold and the genera which have previously been associated with colorectal cancer are surrounded with a box. Abbreviations: FR – fresh-frozen samples; FIT0 – immediately frozen FIT samples; SB0 – immediately frozen stabilization buffer samples.

Next, we identified 21 genera that belong to core microbiome (the number of genera present in over 95% of the samples) (Supplementary table 5). Thereafter, we investigated which out of all significantly different genera belong to the core microbiome and are therefore common in our sample set. In FIT samples, 6 out of 7 significantly different genera belonged to the core. In stabilization buffer samples, 10 out of 16 genera belonged to the core. The significantly different genera belonging to the core are marked bold on Figure 4.

Although 16S sequencing does not provide species level annotation, we compared the genera of the species previously known to be associated with colorectal cancer stages in the multi-population studies from Wirbel et al. 2019 and Yachida et al. 2019, and analyzed if any of the genera differed significantly in our sample types (Supplementary table 6)(14,15). Out of all the taxa significantly differing in FIT0 samples compared to fresh frozen samples, none was shown to be cancer-related (Figure 4). In SB0 samples cancer-related genera such as *Bacteroides* and *Dorea* were significantly different compared to fresh frozen samples (Figure 4). We did not detect *Fusobacteria* which is often associated with colorectal cancer among the genera that are present in 10% of the samples, indicating that the genera is not common among healthy Estonian individuals.

### Gut microbiome composition stability over time

Next, we analyzed if the microbiome composition remains stable in the FIT tubes and stabilization buffer after keeping the samples at room temperature for 4 or 7 days (Figure 1). To evaluate this, FIT0 samples were compared to FIT4 and FIT7 samples and the same was done for SB0, SB4 and SB7 samples. Alpha diversity, beta diversity and differential abundance of the genera were analyzed. We detected no significant differences between FIT samples frozen on different days in terms of the number of observed genera (FDR_FIT0-FIT4_ = 0.2; FDR_FIT0-FIT7_ = 0.9) (Figure 5A) and in Shannon diversity (FDR_FIT0-FIT4_ = 0.099; FDR_FIT0-FIT7_ = 0.12) (Figure 5B) (Supplementary table 4). *Romboutsia* was the only genera with significantly different abundance in samples frozen in day 4 and day 7 compared to immediately frozen FIT samples (FDR < 0.05) (Supplementary Table 7).

**Figure 5.**
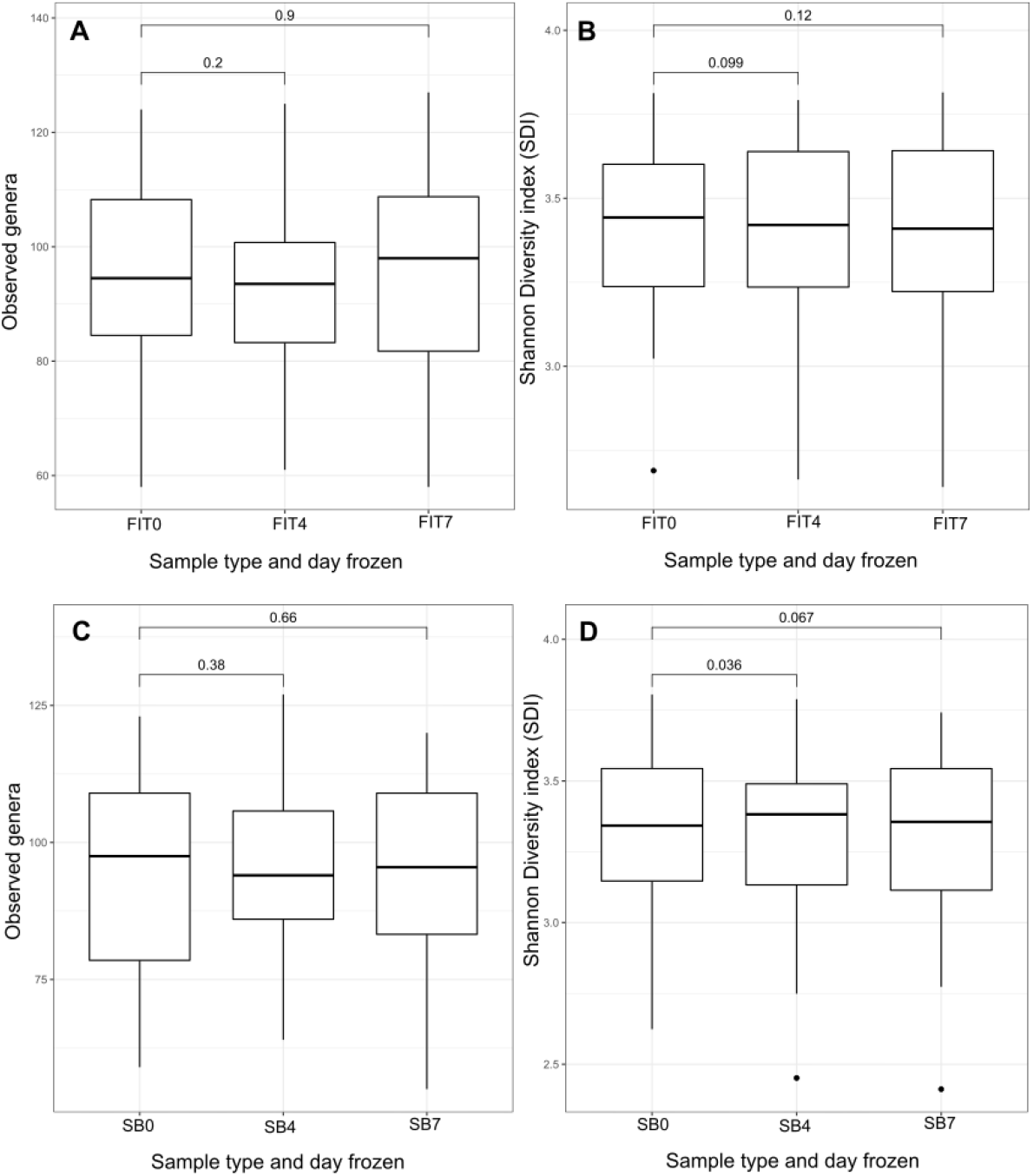
Microbiome diversity between different time-points across sample types. Boxplots represent observed genera (A) and Shannon diversity index (B) in FIT samples and observed genera (C) and Shannon diversity index (D) in stabilization buffer samples from different time points. Abbreviations: FIT0 – immediately frozen FIT samples; FIT4 – FIT samples frozen on day 4; FIT7 – FIT samples frozen on day 7; SB0 – immediately frozen stabilization buffer samples; SB4 – stabilization buffer samples frozen on day 4; SB7 – stabilization buffer samples frozen on day 7.

Similarly, no significant differences were detected in the number of observed genera between SB0 and SB4 or SB7 samples (FDR_SB0-SB4_= 0.38; FDR_SB0-SB7_= 0.66) (Figure 5C), however Shannon index was significantly lower in SB0 compared to SB4 (FDR_SB0-SB4_= 0.036; FDR_SB0-SB7_= 0.067) (Figure 5D) (Supplementary table 4). No genera were significantly different between SB0 and SB4 or SB7 samples (Supplementary Table 7).

When we used all samples from all the time-points for visualizing beta diversity, we observed that the samples remained clustered based on the individual, indicating that the inter-individual differences are bigger compared to storage conditions and days when the sample was frozen (Supplementary Figure 2). This was also supported by PERMANOVA, as the day when the sample was frozen was not significant (R^2^ = 0.0012, p > 0.05). Furthermore, the results were insignificant when the interaction between sample type and day frozen was taken into account (R^2^ = 0.0011, p > 0.05).

## DISCUSSION

Colorectal cancer screening programs all over Europe are using fecal immunochemical test (FIT) as a first step to detect colorectal cancer. As colorectal cancer associated microbiome signatures have been obtained from fresh frozen stool samples from patients with different stages of cancer (14,15), using microbiome for improving colorectal cancer screening has become a topic of high interest. In the current study, we aimed to test whether the FIT tubes used in national CRC screening programs are also suitable for microbiome analysis. The particular FIT tube (QuikRead iFOBT tube) presented in this study has not been previously used to detect microbiome from human stool. We compared the microbial communities detected from FIT tubes with the fresh frozen samples. We also used samples collected in stabilization buffer (DNA/RNA Shield) as an additional collection method for comparison, as stabilization buffers are often used when the collection of fresh-frozen samples is not feasible. Additionally, we evaluated whether the genera that are significantly different in either of the collection methods have been previously associated with colorectal cancer in the multi cohort population studies. Finally, the stability of the microbiome profile for 4 and 7 days was studied in both FIT and stabilization buffer.

Our results indicate that the microbial communities obtained from fresh-frozen samples and FIT tubes are highly similar. Analysis of microbial alpha-diversity demonstrates that the number of genera were not statistically different between the storage methods. Small differences were identified in the Shannon index with microbiome, proving FIT to be less diverse when compared to fresh-frozen samples. However, beta-diversity analysis clearly showed that the differences between subjects were greater than differences between different storage methods. This is in accordance with previous studies that have found interindividual differences to be greater than intraindividual differences between different collection methods (17,34). This was further confirmed by the results of analysis of variance which also indicated that the storage conditions have minimal effect (~2%) on the composition of gut microbiome. Donor-specific factors like diet, age, medications, stool consistency and host genetics are all the likely underlying cause for the substantial inter-individual beta diversity differences.

When analyzing changes in genus abundances, we found that FIT tubes capture a similar community to fresh frozen samples as only 4% of the genera were significantly different between the two collection methods. Even though the most of the significantly different genera belonged to the groups found in the core microbiome (i.e. present in 95% of the samples), none of them have been previously associated with colorectal cancer in multi-cohort population studies (14,15). This supports indicating the possibility to use the FIT tubes for studying the microbial biomarkers related to colorectal cancer.

Next, we wanted to see if the stability of the microbial community could be affected by longer shipping as the time after collecting the initial fecal sample and sending it to the study center for occult blood testing can take up to a week. When analyzing the effect of the storage time, the analysis of variance indicated that the effect of the day when the samples were frozen was not significant. Furthermore, upon comparing the microbial community of FIT samples with different storage times to immediately frozen FIT tubes, no differences in alpha diversity values were detected. Again, beta diversity analysis illustrated that the inter-individual differences were greater than intra-individual differences when data about day 4 and day 7 samples were compared to the samples which were frozen immediately. Differential abundance analysis revealed only one genus (*Romboutsia*) to be significantly different in day 4 and day 7 samples compared to immediately frozen FIT samples, indicating that the abundance of the vast majority of the genera is stable at least for a week. This is in accordance with previous studies which show that FIT tubes can preserve gut microbiome from moderate to excellent levels (4,17,34) and the collection method and time at ambient temperature explain only low amounts of variability (< 10%) (17). These studies have, however, all used FIT tubes from other manufacturers (OC-Sensor and OC-Auto^®^ FIT).

Our study also included a stabilization buffer sample. Previous studies indicate that using a stabilizing buffer is necessary when samples cannot be frozen immediately as certain bacterial taxa start to bloom in untreated samples after spending days in room temperature (17,35). We found that the microbiome community in the stabilization buffer is slightly less similar to fresh frozen samples compared to FIT tubes. Although the community was relatively similar to fresh-frozen tubes in terms of microbial diversity and remained stable up to 7 days, the number of significant differences were higher in colorectal-cancer related bacteria, core genera as well as in the number of significantly different genera compared to the differences found between FIT and fresh frozen samples. Additionally, Shannon diversity was significantly lower compared to fresh frozen samples as well as when comparing SB0 to SB4 samples.

In summary, our results show that FIT tubes are suitable for storing fecal samples for microbiome studies as the captured microbiome profile is similar to fresh frozen samples and remains stable up to 7 days. However, the actual ability to detect cancer specific bacterial signatures with this collection method needs to be confirmed in the future using specific phenotypes such as FIT positive patients with and without colorectal cancer among other diseases. Analyzing the CRC-specific microbiome profile from the same FIT tubes used for fecal occult blood testing would allow to improve the detection of CRC with additional microbiome-based biomarkers. This could potentially make the CRC diagnostics more sensitive and cost-efficient. Fecal samples from CRC screening programs can provide a great resource for biomarker discovery and possibly lead to earlier detection of cancer or prior to its onset (pre-cancer state). In addition, FIT tubes could also be used in studying the role of microbiome in other diseases like pancreatic cancer, inflammatory bowel syndrome, and diverticulitis as well as in population studies where samples are often sent via post and using fresh-frozen samples is not possible. Future studies should also investigate the possibility to use the stool samples collected in FIT tubes for metagenomics and metabolomics analysis, which could provide additional information for early CRC detection.

## Acknowledgements

We thank all the participants for participating in this study and University of Tartu core facility for sequencing the samples. The authors acknowledge the financial support provided by COST-European Cooperation in Science and Technology to the CA17118 “Identifying Biomarers Through Translational Research for Prevention and Stratifiication of Colorectal Cancer.

---

**Supplementary Figure 1.**
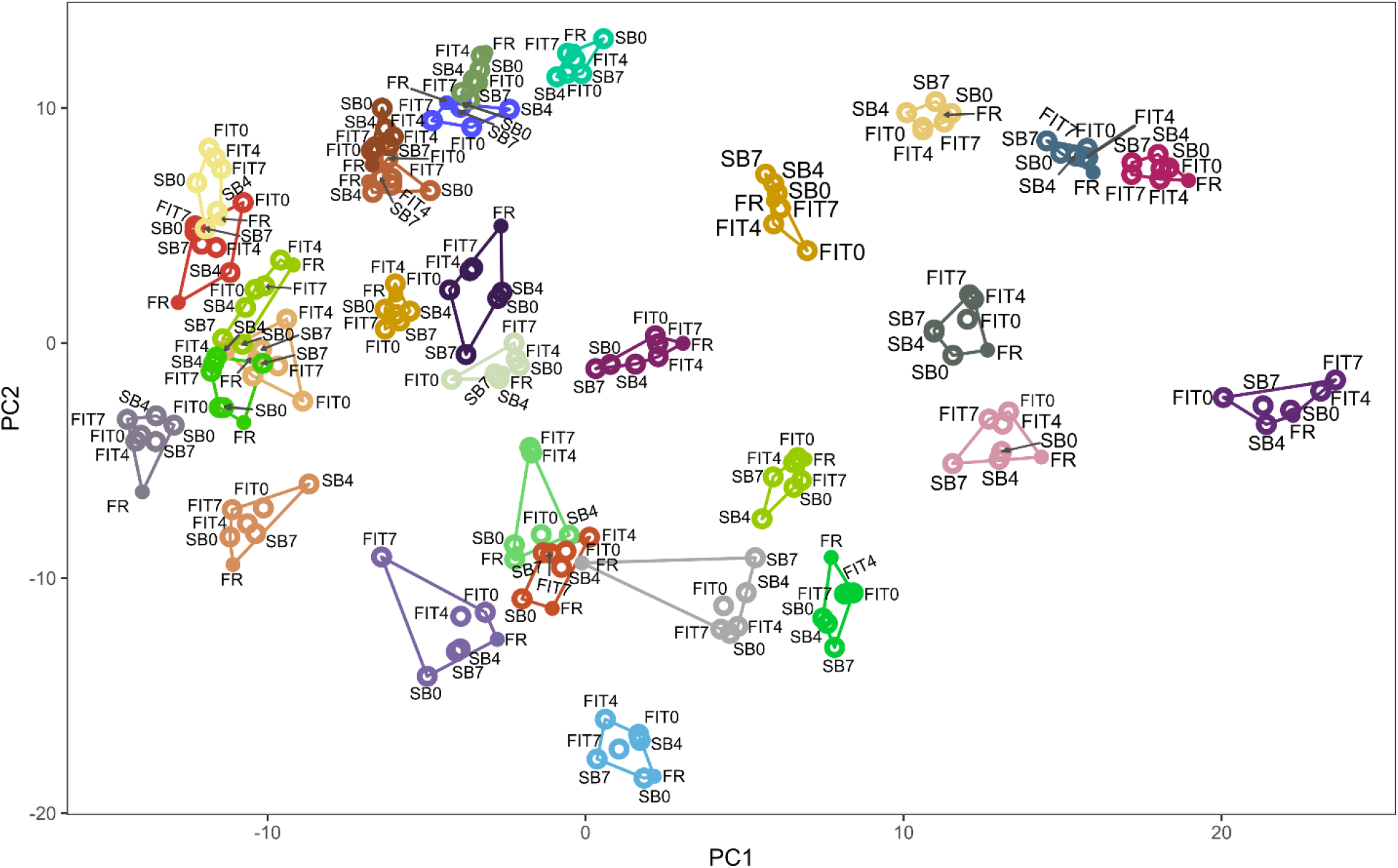
PCA plot showing the differences in the samples collected using different storage conditions including all samples taken from the volunteers. Samples are colored and linked based on the Patient ID. Abbreviations: FR – fresh-frozen samples; FIT0 – immediately frozen FIT samples; FIT4 – FIT samples frozen on day 4; FIT7 – FIT samples frozen on day 7; SB0 – immediately frozen stabilization buffer samples; SB4 – stabilization buffer samples frozen on day 4; SB7 – stabilization buffer samples frozen on day 7.

